# Structures of *Bdellovibrio bacteriovorus* phosphoglucose isomerase reveal novel rigidity in the active site of a selected subset of enzymes upon substrate binding

**DOI:** 10.1101/2021.03.26.437203

**Authors:** R.W. Meek, I.T. Cadby, A.L. Lovering

## Abstract

Glycolysis and gluconeogenesis are central pathways of metabolism across all domains of life. A prominent enzyme in these pathways is phosphoglucose isomerase (PGI) which mediates the interconversion of glucose-6-phosphate and fructose-6-phosphate (F6P). The predatory bacterium *Bdellovibrio bacteriovorus* leads a complex lifecycle, switching between intraperiplasmic replicative and extracellular “hunter” attack-phase stages. Passage through this complex lifecycle involves different metabolic states. Here we present the unliganded and substrate bound structures of the *Bdellovibrio bacteriovorus* PGI, solved to 1.74Å and 1.67 Å, respectively. These structures reveal that an induced-fit conformational change within the active site is not a pre-requisite for the binding of substrates in some PGIs. Crucially, we suggest a phenylalanine residue, conserved across most PGI enzymes but substituted for a glycine in *Bdellovibrio* and other select organisms, is central to the induced-fit mode of substrate recognition for PGIs. This enzyme also represents the smallest conventional PGI characterised to date and likely represents the minimal requirements for a functional PGI.

## Introduction

*Bdellovibrio bacteriovorus* (*B. bacteriovorus*) is a small predatory deltaproteobacterium that preys on other Gram-negative bacteria. *B. bacteriovorus* leads a complex bi-phasic lifecycle switching between extracellular predatory and intraperiplasmic replicative phenotypes; using swimming and/or gliding motility, *B. bacteriovorus* seeks out prey and, upon encounter, attaches to and invades the periplasm of the prey cell. Within the periplasm *B. bacteriovorus* remodels the prey cell into a transient structure called the bdelloplast, wherein it progressively metabolises the cytoplasm and periplasm of its host and uses the nutrients it harvests to grow and divide into new progeny cells. Upon exhaustion of nutrients, *B. bacteriovorus* progeny lyse their host cell and exit in search of new prey cells to repeat the lifecycle. With the sharp rise of antibiotic resistance, *B. bacteriovorus* has shown promise and is currently being investigated as a “living antibiotic” option for treating multidrug resistant Gram-negative bacterial pathogens^1^.

Prior studies have indicated that *B. bacteriovorus* is unlikely to utilise polysaccharides as the primary substrates for energy metabolism and instead respires amino acids to provide energy needed for intraperiplasmic growth^2^. In agreement, high activities of citric acid cycle enzymes and low activities of glycolytic enzymes are observed in *B. bacteriovorus* cell extracts, with the notable exceptions of phosphoglucose isomerase (PGI) and glyceraldehyde 3-phosphate dehydrogenase^3^. Sequencing of the *B. bacteriovorus* genome has revealed a full complement of enzymes required for production of ATP through glycolysis^4^. Why high *B. bacteriovorus* PGI (*Bb*PGI) activity is detected in *B. bacteriovorus* cell extracts, when glycolysis is not the primarily route for energy production, is unknown^3^.

PGI mediates reversible isomerization of glucose-6-phosphate (G6P) to fructose-6-phosphate (F6P) in glycolysis and gluconeogenesis. The catalytic mechanism of PGI mediated isomerization has been well documented using mammalian PGIs as model systems ^5-9^. The process begins with the PGI opening the substrate ring before mediating a proton transfer, using a *cis-*enediol mechanism, between the C1 and C2 position before ring closure. The mechanism of bacterial PGIs is assumed to be identical, owing to bacteria having analogous structures with similar domain arrangements. However, a relatively small number of bacterial PGIs have been structurally characterised, and these represent only a small fraction of total prokaryotic classes. In particular, no structure is available from a deltaproteobacterial source.

In addition to the classical role of isomerisation, PGIs often function as ‘moonlighting’ proteins. In humans, PGI moonlights as autocrine motility factor, neuroleukin, and differentiation and maturation mediator^10–12^. Conversely, in *Lactobacillius crispatus*, PGI moonlights as a pH-dependent adhesion protein^13^. Thus, PGI enzymes can have diverse functions in addition to their roles in central metabolism, and there are likely additional roles for this family yet to be discovered. *Bb*PGI is part of a novel grouping of smaller PGIs which are typically ~100 amino acids (aa) smaller than other conventional PGIs (for example, *Escherichia coli* PGI is 549 aa compared to only 408 aa for *Bb*PGI). All structurally characterised PGIs reside in one of two groupings, those above 540 aa and those below 450 aa in size. To gain valuable insights into the role of *Bb*PGI, we have obtained two crystal forms of *Bb*PGI which we have solved to either 1.74 Å and 1.84 Å. Using ligand soaking experiments, we have obtained a crystal structure of *Bb*PGI bound to its substrate. Through these crystal structures we determine that the *Bb*PGI active site, in contrast to those of previously characterised PGIs, does not undergo significant conformational change upon substrate binding. This lack of active site mobility can be partially attributed to the substitution of a Phe residue, conserved in most PGIs, for a Gly residue in *Bb*PGI. We also note a novel feature in the smaller PGIs wherein the hook region is in a different position to that of the larger PGI structures.

## Results

### Overall structure of *Bb*PGI

To determine the *Bb*PGI structure we used full-length recombinant *Bb*PGI protein in crystallography experiments. Crystallization trials yielded two crystal forms of *Bb*PGI which were processed in P12_1_1 and P3_1_21 spacegroups (1.74 Å and 1.84 Å, respectively). Data from the P3_1_21 form was solved by molecular replacement using PDB 1B0Z from *Geobacillus stearothermophilus* as a search model^14^. Data processing and refinement statistics are displayed in Table 1. The P3_1_21 form structure was used as a search model for molecular replacement solution of the P12_1_1 dataset. The P3_1_21 form had a single copy of *Bb*PGI comprising the asymmetric unit, whereas the P12_1_1 dataset asymmetric unit is composed of 8 copies. The eight copies in the P12_1_1 dataset form four largely identical dimers (RMSDs of 0.107-0.190 Å between dimers). A dimeric assembly also exists in the P3121 form but is generated via crystallographic symmetry axes. Dimer partners are related to one another by a 2-fold rotational symmetry. The polypeptide backbone for residues 3-406 can be fully traced into the electron density map, with only the first and last two amino acids of the polypeptide sequence and the His-tag being disordered and absent from the final refined models.

**Table.1.** Data collection and refinement statistics.

|  | *Bb*PGI crystal form 1 | *Bb*PGI crystal form 2 | Ligand bound *Bb*PGI |
| --- | --- | --- | --- |
| Accession code | 7NSS | 7O0A | 7NTG |
| **Data Collection** |  |  |  |
| Spacegroup | P3<sub>1</sub>21 | P12<sub>1</sub>1 | P3<sub>1</sub>21 |
| Cell dimensions |  |  |  |
| a, b, c (Å) | 101, 101, 76.7 | 137.9, 118.1, 139 | 101.2, 101.2, 76.4 |
| α, β, γ (°) | 90, 90, 120 | 90, 118, 90 | 90, 90, 120 |
| Resolution | 57.68 (1.84) | 59.54 (1.74) | 43.82 (1.67) |
| R<sub>merge</sub> | 0.101 (1.919) | 0.090 (0.607) | 0.057 (1.966) |
| R<sub>p.i.m</sub> | 0.034 (0.821) | 0.073 (0.495) | 0.019 (0.655) |
| I/σI | 23.2 (1.3) | 8.7 (1.5) | 25 (1.5) |
| CC<sub>1/2</sub> | 1 (0.559) | 0.982 (0.615) | 1 (0.822) |
| Completeness | 95.8 (99.9) | 98.8 (99.1) | 99.9 (99.9) |
| Multiplicity | 18.7 (12.5) | 3.9 (3.4) | 19.6 (19.4) |
| **Refinement** |  |  |  |
| Resolution | 1.84 | 1.74 | 1.65 |
| No. of reflections | 37892 | 375276 | 52622 |
| R<sub>work</sub>/R<sub>free</sub> | 0.184/0.221 | 0.222/ 0.253 | 0.178 / 0.217 |
| No. of atoms |  |  |  |
| Protein | 3191 | 26060 | 3217 |
| Ligand | 4 | 24 | 20 |
| Water | 249 | 3254 | 283 |
| **B-factors** |  |  |  |
| Protein | 36.7 | 21.8 | 37.9 |
| Ligand | 50.8 | 30.7 | 40.5 |
| Water | 40.8 | 30.9 | 43.8 |
| R.m.s. deviations |  |  |  |
| Bond lengths | 0.0146 | 0.0135 | 0.0153 |
| Bond angles | 1.77 | 1.68 | 1.83 |

The core structure of the *Bb*PGI is similar to other characterized PGI structures but lacks much of the loop decoration characteristic of larger GPIs (**Figure 1**). *Bb*PGI is comprised of three domains: the large domain, consisting of residues 1-31 and 213-384 which forms a mixed 5-strand β-sheet sandwiched by α-helices; the small domain, consisting of residues 44-195 which forms a parallel 5- strand β-sheet flanked by α-helices; and the C-terminal arm sub-domain, a single α-helix comprised of residues 392-408 (**Figure 1A**). The small and large domain are linked to each other by residues 32-43 and 196-212, with residues 199-211 forming an α-helix. A small loop (residues 385-391) connects the large domain to the C-terminal arm subdomain. A hook region (residues 306-348) extends from the large domain and packs against the small domain of the dimer partner providing dimer-stabilising hydrogen bonding and hydrophobic interactions (**Figure 1B**). We note that positioning of the hook region varies in PGIs, with the interaction loop extending from either the N-terminus or C-terminus of α20 (*Bb*PGI helix numbering) to interact with either the small domain or large domain of the dimer partner, respectively (**Figure S1**). In PGIs of size >540 aa the loop extends from the C-terminal position to contact the large domain (determined using 15 unique representative structures in the PDB). Conversely, in *Bb*PGI and PGIs of <450 aa, an alternative conformation is adopted with extension of the loop from the N-terminus of α20 (6 unique representative structures). This is an intuitive observation since α20 tilts away from the dimer partner, with the N-terminus being much closer for contact to the dimer partner than the C-terminus, thus requiring less residues to make dimer contact. The C-terminal arm sub-domain in *Bb*PGI sandwiches the hook region between itself and the small domain, whereas in larger PGIs the hinge region wraps around the outside of the C-terminal arm sub-domain.

**Figure 1.**
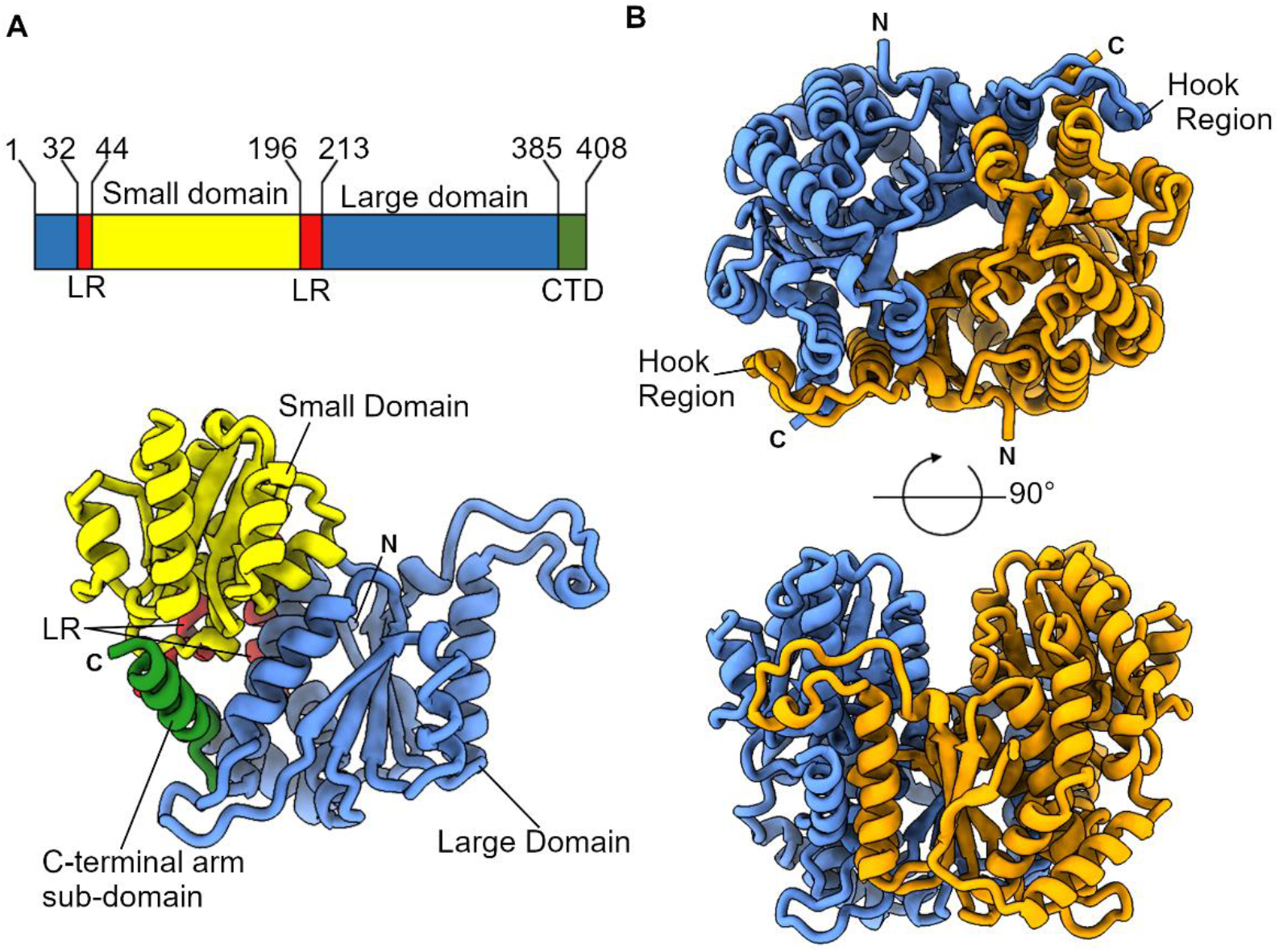
Structure of *Bb*PGI. A, Domain boundaries and their corresponding position in *Bb*PGI. LR, Linker regions; CTD, C-terminal domain. B, Orthogonal views of the *Bb*PGI dimer assembly with dimer partners coloured blue and orange. The hook region which contributes to the dimeric arrangement is labelled. C-terminus and N-terminus are marked by C and N, respectively.

All three domains contribute to the proposed active site, with the small domain positioned to coordinate the phosphoryl group of the incoming substrate. Active site architecture is completed by recruitment of the large domain of a dimer partner. This is necessary for catalysis since the dimer partner donates the conserved residue H285 required for sugar ring opening/closing. *Bb*PGI is one of the smallest, conventional PGI family enzymes reported to date, with DALI analysis revealing strong structural homology to PGIs from the thermophiles *Thermotoga maritima* (PDB 2Q8N, Z-score: 48.6) and *Thermus thermophilus* HB8 (PDB 1ZZG, Z-score: 48.2)^15^. The *Bb*PGI dimeric assembly is supported by a PISA calculated interface area of 4811 Å^2^ (25% of total monomer surface area) between dimer partners, suggesting the dimer to be physiologically relevant^16^. However, the *Thermus thermophilus* HB8 PGI exists in solution as a dynamic equilibrium between monomer and dimer and, owing to the high structural homology, *Bb*PGI may also exist in a similar equilibrium state^15^.

### The *Bb*PGI active site and ligand accommodation/coordination

Although the conformation of the active site is similar to larger PGIs we decided to validate substrate binding in this smaller grouping of compact PGIs. To do so, we performed a ligand soaking experiment with G6P using the P3121 crystal form of *Bb*PGI. Strong Fo-Fc difference density was observed in the active site and could be unambiguously modelled as an extended, open ring phosphosugar (**Figure 2**). Since G6P can be interconverted to F6P we queried which had bound to the active site; unfortunately electron density is most diffuse at the C1 position and it is difficult to conclusively state whether a tetrahedral (G6P) or trigonal planar (F6P) geometry is present at the C2 position. However, the F6P O2 can be more-confidently modelled into the electron density than the G6P O2, thus we have tentatively modelled F6P into the active site and will refer to it as such hereafter. It is possible this ambiguity is the result of a mixed population of open-ring G6P and F6P being bound.

**Figure 2.**
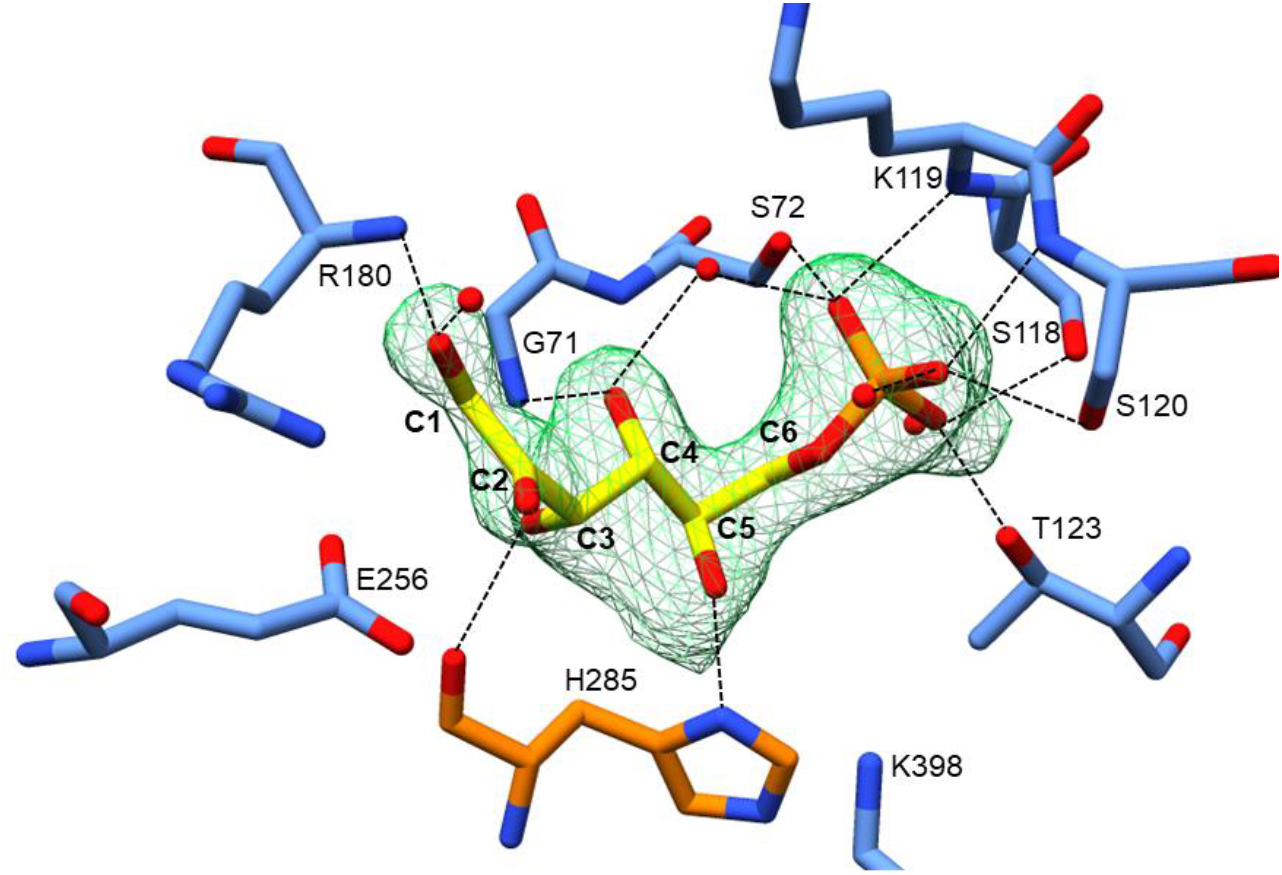
The coordination of F6P in the active site. A F_o_-F_c_ omit map indicating the presence of F6P (yellow) is depicted as a green mesh (contoured at 3 σ). The ligand is coordinated by numerous hydrogen bonds represented by dashed lines. Peptide backbone displayed as sticks with opposing protomers of the dimer coloured in blue and orange.

In the active site, the phosphoryl group of F6P is extensively coordinated by the small domain through the hydroxyl-containing sidechains of S72, S118, S120 and T123, the backbone nitrogen of K119 and S120, and three ordered waters (**Figure 2**). F6P is also coordinated through O1 interacting with the backbone nitrogen of R180 (small domain) and an ordered water. F6P interactions are further provided by: O3 bonding with the backbone nitrogen of H285, O4 binding the backbone nitrogen of G71 and an ordered water, and O5 interacting with the sidechain of H285 (**Figure 2**). Active site residues known to be involved in mediating isomerisation (R180, E256, H285 and K398) are positioned and orientated similarly to other PGIs (**Figure 3**).

**Figure 3.**
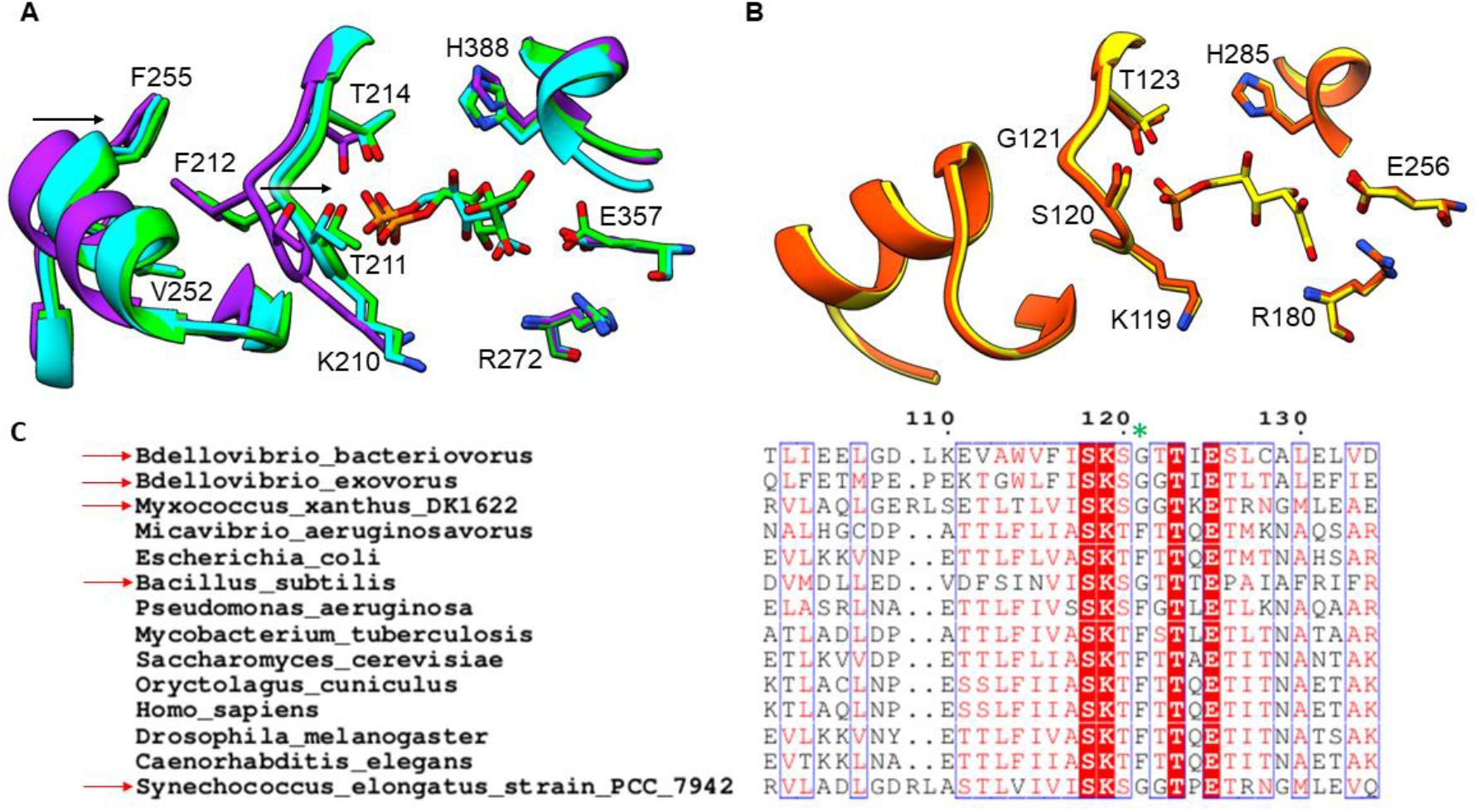
Tracking conformational changes in the active site. A, rPGI (unliganded, purple) loops 209-215 and 245-259 move towards the active site upon F6P (green) or 5-phospho-D-arabinonate (cyan) binding to interact with the phosphoryl group (indicated by arrow). Binding of ligand also drives residues 385-389 towards the active site to re-position the catalytic histidine (H388). B, Minimal backbone conformational movements between unliganded (orange) and ligand bound (cyan) *Bb*PGI. C, Sequence alignment of *Bb*PGI against PGIs of close relatives and model organisms. Numbered relative to *Bb*PGI with red arrows indicating organisms with a Gly at the semi-conserved Phe position (green asterisk).

### Conformational changes at the *Bb*PGI active site

Upon ligand binding in other PGIs, relatively-mobile active site residues facilitate hydrogen bonding with the phosphoryl group of G6P or F6P. This is evident in the unliganded, F6P and 5-phospho-D-arabinonate bound structures of rPGI (PDBs 1HM5, 1HOX and 1G98, respectively)^5–7^, wherein loops 209-215 and 245-259 move towards the active site upon ligand binding (**Figure 3A**)^7^. This movement transitions the protein from an “open” to a “closed” state. The synchronised movement of these loops has been attributed to F212, which connects the 209-215 loop to the 245-259 loop through a hydrophobic interaction network^7^. Interestingly this induced-fit mechanism is not witnessed in our two states of *Bb*PGI and the loop arrangement between unliganded and ligand bound is identical, with the enzyme held in the “closed” state regardless of ligand occupancy (**Figure 3B**). The equivalent residue to rPGI F212 is G121 in *Bb*PGI and does not connect the two loops. Although heavily conserved, this Phe residue (rPGI F212) is sometimes switched to a Gly in some organisms (**Figure 3C**). Notably use of a Gly residue is apparent in various bacteria, including close relatives of *Bdellovibrio.* This could indicate that these species undergo smaller or no conformational changes in this region upon ligand binding, since the 245-259 loop may not be connected to the equivalent 209-215 loop (rPGI numbering). Interestingly, all PGI structures <450 aa utilize a Gly, while in PGI structures of >540 aa this residue is a Phe. A second conformational change is observed in rPGI wherein upon F6P binding, wherein residues 385-389 are also recruited to the active site repositioning H388 for catalysis (**Figure 3A**)^7^. Our structure undergoes no conformational changes in the equivalent histidine residue (H285) upon ligand binding. Previous work using cis-enediol(ate) intermediate mimics in rPGI noted a conformational change in residues 513-521 which brought K518 towards the bound ligand. Again, we observe no conformational changes in the equivalent residues (393-401) upon F6P binding, agreeing with previous G6P complex structures^9, 17^. It is possible only cis-enediol(ate) intermediates drive this particular conformational change.

## Discussion

In solution, G6P and F6P exist primarily in cyclic hemiacetal and hemiketal forms^18^. Our structure presents F6P bound in an open ring conformation strongly suggesting the enzyme has facilitated ring opening. In addition, placement of active site residues is consistent with other active PGIs. It is interesting that *Bb*PGI undergoes no conformational changes upon ligand binding unlike other characterised PGIs. Certainly, it is apparent that conformational movement in the polypeptide backbone is not a prerequisite for isomerase activity, which may also be the case in other, similar PGIs. The *Leishmania mexicana* eukaryotic parasite *PGI* has also been demonstrated to undergo no ligand driven conformational changes, however unlike our structure, it is held in the “open” conformation rather than the “closed” state (the authors suggest the loop would still be required to move ~5 Å during isomerisation in order to be within a suitable distance of the catalytic glutamate residue)^19^. The positioning of a Gly (G121) instead of a Phe residue in the active site may also be indicative of PGIs which do not reposition loops for binding of the phosphoryl group. The structure of *Bb*PGI (408 amino acids) is significantly smaller than other bacterial/mammalian PGIs (normally >500 amino acids) and matches PGIs from thermophiles, which likely need to be more rigid/compact to function at high temperatures. Certainly, this structure is likely to approximate the minimum structural requirements for an active PGI. It would be interesting if the compact nature of *Bb*PGI was an adaptation particular to the predatory lifestyle of *B. bacteriovorus.* The reasoning as to why other PGIs have not evolved to be more compact and still have ligand induced conformational changes is unknown and may relate to their active sites/states being involved in other moonlighting functions. From our structures it is not obvious why PGI activity is noticeably higher than other glycolytic processes in *B. bacteriovorus* cell extracts.

We have observed that the open reading frame of *Bbpgi* is separated by only 38 nucleotides from the diguanylate cyclase *dgcb*, although they are unlikely to share an operon (personal communication with Luke Ray in R.E Sockett lab). Deletion of *dgcb* abolishes the ability of *B. bacteriovorus* to initiate host cell invasion^20^. We have previously characterised DgcB as a diguanylate cyclase likely activated by a phosphorylation event, which drives production of a localised pool of cyclic-di-GMP that signals to downstream proteins to initiate invasion^21^. Remarkably, enzymes from metabolic pathways have been isolated from *B. bacteriovorus* through a c-di-GMP capturing compound^22, 23^. Additionally, c-di-AMP has also been implicated in binding metabolic enzymes, and G6P has recently been directly correlated to regulating c-di-GMP levels^24, 25^. Considering the propensity of PGI enzymes to take on moonlighting roles we wondered whether *Bb*PGI links c-di-GMP signalling to metabolism by directly interacting with DgcB. We carried out pulldown assays to test this hypothesis, however results indicated that they did not interact under the conditions assayed (**Figure S2**). We cannot, however, exclude the possibility that the wildtype variants of DgcB (c-di-GMP bound dimer) and/or *Bb*PGI exist in a state non-permissive for interaction or require an additional third protein complexed to facilitate interaction, or that G6P/F6P regulate c-di-GMP signalling through an unknown mechanism.

In summary, *Bb*PGI is representative of the minimal PGI unit and highlights that, through substitution of the Phe residue found in most PGIs (rPGI F212) to a Gly, a less drastic or absent conformational change in the active site is likely to occur in enzymes with this variant. Whether this lack of mobility impacts upon PGI catalysis and its ability to interact with other ligands and protein partners remains an open question.

## Methods

### Cloning and protein production

The full open reading frame for *Bbpgi* (UNIPROT entry Q6MPU9, gene *bd0741*, amino acids 1-408) was amplified from *B. bacteriovorus* HD100 genomic DNA using primers detailed in **Table S1** Amplified construct DNA was inserted into a modified pET41c plasmid (Novagen, GST-tag removed and thrombin cleavable 8xHis-tag introduced at C-terminus) by restriction free cloning. Correct plasmid construct was confirmed by sequencing. Plasmid was transformed into *Escherichia coli* expression strain BL21λDE3 (New England Biolabs) for expression. Cells were grown in a shaking incubator (180 RPM) at 37 °C in 2 x LB supplemented with 100 μg/ml Kanamycin until an OD600 of 0.6-1 was reached. Expression was induced with 0.5 mM IPTG before 15 °C overnight shaking incubation. Cells were harvested by centrifugation at 6675 × g for 6 min (4 °C) before being frozen at −20 °C.

## Protein purification

Cell pellets were defrosted and resuspended in Buffer A (20 mM Imidazole pH 7.0, 400 mM NaCl and 0.05% Tween 20) with lysozyme. Cells were lysed on ice by sonication and lysate was clarified by centrifugation at 48,400 g for 1 hour (4 °C). Supernatant was loaded onto buffer A equilibrated HisTrap FF nickel columns (GE Healthcare). Columns were washed with 12 CVs of buffer A before two 20 ml elution steps at 8% buffer B (400 mM Imidazole pH 7.0, 400 mM NaCl and 0.05% Tween 20) in buffer A and 100 % buffer B. The purity of samples was confirmed by SDS-PAGE. Eluted protein was dialysed overnight at 4 °C against 20 mM HEPES pH 7.0 and 200 mM NaCl. Protein was concentrated to 40 mg/ml using Vivaspin^®^ spin-concentrators (Sartorius) and snap-frozen.

## Crystallization and structure determination

Protein at 40 mg/ml was screened for conditions to promote crystallization (1:1 ratio of protein to reservoir solution, 4 μl total drop size). Diffraction quality crystals were obtained in 0.2 M ammonium acetate, 0.1 M sodium acetate pH 5.0 and 20% PEG 4000 (crystal form 1), and 0.2 M ammonium acetate, 0.1 M sodium acetate pH 4.0 and 15% PEG 4000 (crystal form 2). Crystals were cryoprotected in mother liquor supplemented with 20% ethylene glycol (v/v) before being flash cooled in liquid nitrogen. To obtain the F6P bound *Bb*PGI structure, crystal form 1 was soaked in mother liquor supplemented with 3 mM G6P and 20% ethylene glycol for 5 min before being flash cooled in liquid nitrogen. Diffraction data were collected at Diamond Light Source in Oxford, UK. Data reduction and processing was performed through the *xia2* suite^26^. Crystal form 1 (P3121 spacegroup) was successfully phased by molecular replacement in MRage using PDB accession 1B0Z as a search model^27^. Phenix autobuild was used to partially rebuild incorrect sections of the model^28^. Crystal form 2 (P1211 spacegroup) and the G6P soaked crystal form 1 data were solved by MRage molecular replacement using crystal form 1 as a search model. Models were finalised through iterative manually building in *Coot* and refinement in REFMAC^29, 30^. Structure representations were produced in Chimera and ChimeraX^31, 32^.

## Sequence analysis

Protein sequences were aligned through Clustal Omega and figures produced through ESPript3^33, 34^. UNIPROT identifiers used: Q6MPU9 (*B. bacteriovorus)*, M4VS33 (*Bdellovibrio exovorus*), Q1CX51 (*Myxococcus xanthus* DK1622), G2KQS5 (*Micavibrio aeruginosavorus*), Q1R3R3 (*Escherichia coli*), P80860 (*Bacillus subtilis*), Q02FU0 (*Pseudomonas aeruginosa*), P9WN68 (*Mycobacterium tuberculosis*), P12709 (*Saccharomyces cerevisiae*), Q9N1E2 (Rabbit; *Oryctolagus cuniculus*), P06744 (*Homo sapiens*), P52029 (*Drosophila melanogaster*), Q7K707 (*Caenorhabditis elegans*), Q31LL0 (*Synechococcus elongatus* strain PCC 7942)

## Acknowledgements

R.W.M. was supported by a BBSRC MIBTP studentship. This research was funded by the U.S. Army Research Office and the Defense Advanced Research Projects Agency accomplished under Cooperative Agreement Number W911NF-15-2-0028. The views and conclusions contained in this document are those of the authors and should not be interpreted as representing the official policies, either expressed or implied, of the Army Research Office, DARPA, or the U.S. Government. The U.S. Government is authorized to reproduce and distribute reprints for Government purposes not withstanding any copyright notation hereon. We would like to thank Diamond Light Source for access to beamline I04 (under proposal numbers mx10369 and mx14692)

