## Supplementary Materials for "Structures of *Bdellovibrio bacteriovorus* phosphoglucose isomerase reveal novel rigidity in the active site of a selected subset of enzymes upon substrate binding"

**Supplementary Table.1** Primers used to clone *Bb*PGI into a modified pET-41c vector

| Primer direction | Primer Sequence |
|---|---|
| Forward (5' to 3') | GCAGCGGCCTGGTGCCGCGCGGCAGCCATATGCACGTTATGTTGGAGATTTCGCATTCG |
| Reverse (5' to 3') | CTCAGTGGTGGTGGTGGTGGTGCTCGAGTCAAGCTTTCTTCAATTTCTCTTTCG |

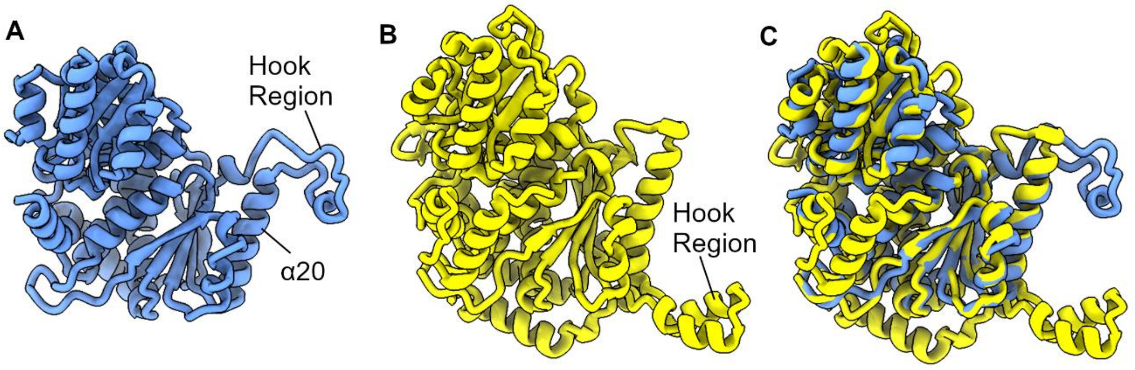


**Supplementary Figure.1**. Positioning of the hook region comparison. A, Structure of *Bb*PGI with hook region N-terminal of α-helix 20. B, Structure of rabbit PGI (PDB code: 1HM5) with hook region C-terminal of equivalent α-helix to *Bb*PGI α-helix 20. C, Superimposition of A onto B demonstrating differences in hook region positioning.

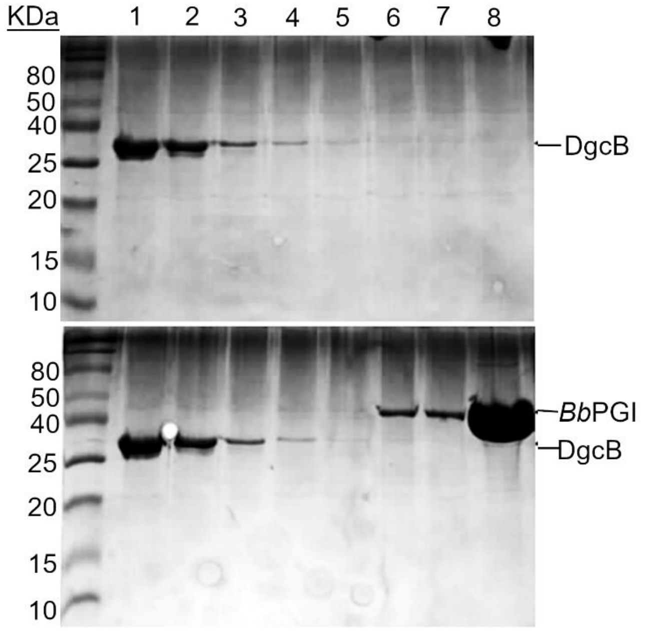


**Supplementary Figure.2.** SDS-PAGE gels of a pulldown assay investigating whether *Bb*PGI interacts with thrombin cleaved wildtype DgcB. Top panel, DgcB only control showing DgcB is unable to bind to the nickel resin. Bottom Panel, *Bb*PGI and DgcB incubated together, no coelution of DgcB with *Bb*PGI observed suggesting no interaction. 1) Flowthrough; 2) Wash1; 3) Wash2; 4) Wash3; 5) Elution 1, 8% buffer B; 6) Elution 2,8% buffer B; 7) Elution 3, 100% buffer B; 8) Elution 4, 100% buffer B.

### Supplementary Methods

#### Pulldown Interaction Assay

DgcB was purified by previously reported protocols[1], and dialysed against 500 mM NaCl, 2.5 mM $CaCl_2$, and 20 mM HEPES pH 8.3 with 4 ul of thrombin (Merck Millipore) for two days at 4 °C. Un-cleaved protein was removed by passing solution over a 5 ml HisTrap FF nickel column (GE Healthcare) before elution with 20 ml of buffer A (from main body methods). Eluted protein was dialysed overnight into 20 mM HEPES pH 7.0 and 200 mM NaCl. To remove thrombin from the sample, dialysed protein was passed over a 1 ml HiTrap Benzamidine FF column (GE Healthcare) and concentrated with Vivaspin® spin-concentrators (Sartorius) to 35.5 mg/ml. Nickel beads (100 µl) were equilibrated in buffer A and transferred to a microcentrifuge tube. Thrombin cleaved DgcB and *Bb*PGI were added to the beads (2 mg total of each). Buffer A was added to the beads to a final volume of 1 ml. Protein mix was rotated overnight at 4 °C to allow proteins to interact. Protein suspension was added to a 0.8 ml Bio-spin chromatography column (Bio-rad) and centrifuged at 300 RPM for 45 secs (flowthrough collected). Beads were washed three times with 500 µl buffer A, before elution with two 500 µl solutions of 8% buffer B in buffer A (v/v) and two 500 µl solutions of 100 % buffer B. All wash and elution steps were by centrifugation (300 RPM for 45 secs) in a benchtop centrifuge. Results were analysed by SDS-PAGE.
